# Effects of aperiodic neural activity on sleep-based emotional memory consolidation across the lifespan

**DOI:** 10.1101/2025.03.30.646183

**Authors:** Zachariah R. Cross, Amanda Santamaria, Scott W. Coussens, Mark J. Kohler

## Abstract

Sleep neurophysiology undergoes significant changes across the lifespan, which coincide with age-related differences in memory, particularly for emotional information. However, the mechanisms that underlie these effects remain poorly understood. One potential mechanism is the aperiodic component, which reflects “neural noise”, differs across age, and is predictive of perceptual and cognitive processes. In this study, we investigated how intrinsic (i.e., resting-state) aperiodic neural activity modulates sleep-based emotional memory consolidation across the human lifespan. In a within-subjects, repeated measures design, forty-two participants aged 7 – 72 years (*M* = 26.60, *SD* = 17.45; 26 female) completed a learning and baseline recognition emotional memory task before a 2hr afternoon sleep opportunity and an equivalent period of wake. Recognition accuracy was also assessed post-delay. We found that aperiodic slopes follow a u-shaped trajectory across the lifespan: slopes flatten from childhood to young adulthood, before steepening thereafter, with this effect most prominent in frontal regions. Age-related differences in aperiodic slopes also explained interindividual differences in emotional memory consolidation, with less age-related flattening of slopes associated with stronger consolidation of negative stimuli post-sleep but not post-wake. Lastly, independent of aperiodic activity, age-related differences in NREM oscillatory activity predicted emotional memory consolidation. These findings suggest that the efficiency of sleep-based emotional memory consolidation is modulated by age-related differences in aperiodic neural and NREM oscillatory activities, providing novel insights into the neurophysiological mechanisms underpinning emotional memory across the lifespan.

## 1. Introduction

Memory consolidation, particularly during sleep, is a fundamental process for the stabilization and integration of new information (Brodt, Inostroza, Niethard, & Born, 2023). This process is modulated by a variety of neural mechanisms, including oscillatory activity, which plays a crucial role in the successful encoding and retrieval of mnemonic information (Marshall, 2020; Düzel, Penny, & Burgess, 2010). While such mechanisms facilitate memory consolidation more broadly, they have been specifically implicated in the preferential processing and consolidation of emotional memories (van der Helm & Walker, 2009). Emotional memory consolidation during sleep is thought to vary across the lifespan, with age-related differences in both the neural processes underlying memory consolidation and the sleep features that support this process. Similarly, recent research has highlighted that aperiodic neural activity, quantified as the aperiodic slope, may be a critical marker of age-related differences in perceptual and cognitive functioning (Cross et al., 2022a, 2022b; Dziego et al., 2023). In younger individuals, steeper aperiodic slopes may be associated with more efficient memory processes (Preston et al., 2025; Voytek et al., 2015), while older adults tend to exhibit flatter slopes, reflecting less efficient neural processing (Merkin et al., 2023; Thuwal et al., 2021). Understanding how these age-related differences in aperiodic activity interact with sleep-based emotional memory consolidation could provide valuable insights into the mechanisms supporting memory across the lifespan.

### 1.1. Sleep-based emotional memory consolidation across the lifespan

Emotional memory is subject to differential processing across the lifespan, and accordingly the relative impact of sleep on emotional memory consolidation may vary by age. In younger adults, there is an established benefit of sleep for consolidation of emotional information over neutral (Hu, Stylos-Allen & Walker, 2006; Payne, Stickgold, Swanberg & Kensinger, 2008; Wagner, Gais & Born, 2001; Wagner, Hallschmid, Rasch & Born, 2006), with effects extending to short sleep durations such as naps (Alger, Chen, & Payne, 2019; Bennion et al., 2016; Cellini, Torre, Stregagno, & Sarlo, 2016; Nishida, Pearsall, Buckner, & Walker, 2009; Payne et al., 2015).

Similarly in children, greater emotion and arousal interact with sleep to improve recognition (Bolinger et al., 2018; Prehn-Kristensen et al., 2009; Prehn-Kristensen et al., 2013). Children and adolescents show higher proportion of slow wave sleep (SWS), slow oscillations (SO) and spindle activity during NREM sleep (Bolinger et al., 2018; Kaestner et al., 2013; Kurth et al., 2010; Kurdzriel, 2018; Nishida et al., 2009; Rasch & Born, 2013), features of sleep associated with emotional memory processing in adults, and which may suggest greater potential for sleep mediated emotional memory consolidation at younger ages. Results for middle-aged to older adults resemble that of young adults, showing sleep-associated consolidation to benefit emotional information over neutral in naps (Alger, et al., 2018) and overnight sleep (Jones, Schultz, Adams, Baran, & Spencer, 2016), with the exception that older adults show a positivity bias, such that positively toned information is preferentially remembered over negative and neutral information (Jones et al., 2016). Older adults also have a diminished capacity for sleep-associated memory consolidation of declarative information, potentially influenced by reductions in SO activity, sleep spindls, and REM sleep (Diekelmann, Wilhelm & Born, 2009; Rasch & Born, 2013).

### 1.2. Age-related differences in aperiodic neural activity and sleep-based emotional memory consolidation

While younger individuals generally exhibit more efficient memory consolidation during sleep, age-related differences in both cognitive and neural mechanisms often lead to declines in memory function in older adults (Sander, Fandakova, Werkle-Bergner, 2021). One such neural marker of age-related differences in information processing capacities that has gained attention in recent years is the aperiodic component of electroencephalography (EEG), quantified as the aperiodic slope (Donoghue et al., 2020). This measure reflects the level of neural “noise” in the brain, likely arising from a combination of factors, including low-pass filtering properties of dendrites (Buzsáki et al., 2012), frequency-dependent current propagation in biological tissues (Bédard & Destexhe, 2009), stochastically driven damped oscillators characterized by varying relaxation rates (Evertz et al., 2022), and/or the balance between excitation and inhibition (E/I) in neuronal populations (Martínez-Cañada et al., 2023; Wiest et al., 2023). The aperiodic slope is not static across the lifespan: in younger individuals, the aperiodic slope tends to be steeper, reflecting a higher level of neural efficiency and more organized, stable brain activity (Cross et al., 2024; Donoghue et al., 2020; Merkin et al., 2023). In contrast, older adults typically exhibit flatter slopes, which may indicate less efficient neural processing and greater fluctuations in spontaneous brain activity (Bornkessel-Schlesewsky et al., 2022; Voytek et al., 2015). This shift in aperiodic activity across the lifespan is thought to influence a variety of cognitive functions, including those critical to memory (Thuwal et al., 2021; Tran et al., 2020; Voytek et al., 2015). From this perspective, intrinsic (i.e., resting-state) aperiodic slopes may be a useful marker of age-related differences in emotional memory consolidation; however, whether this effect interacts with sleep-based memory consolidation processes remains unknown.

Given that sleep plays a crucial role in memory consolidation, particularly negative and positive events, it may be that aperiodic neural activity modulates sleep-based emotional memory consolidation across age. Flatter aperiodic slopes in older adults may reflect less efficient neural processing during sleep (Helfrich, Lendner, & Knight, 2021), impairing the ability to consolidate emotional memories effectively. By contrast, younger individuals with steeper slopes may experience more efficient memory consolidation, resulting in better stabilization of emotional information during sleep. Recent theoretical models support this prediction (Helfrich et al., 2021): while NREM oscillations, such as SOs, sigma activity, and their coupling, facilitates network-level processing via hippocampal–neocortical interactions (Staresina et al., 2016), aperiodic activity facilitates switching from network-level processing to local processing to enable transformation of mnemonic content (Helfrich et al., 2021). From this perspective, age-related variability in NREM oscillatory activity and intrinsic aperiodic activity might jointly explain age-related differences in (emotional) memory by modulating the balance between network-level processing and local processing during sleep. In younger adults, steeper aperiodic slopes may promote more efficient switching between these processes, facilitating the stabilization of emotional memories via the coupling of NREM oscillations (such as SOs and sigma activity) with hippocampal– neocortical interactions. In contrast, flatter aperiodic slopes in older adults may disrupt this balance, impairing the transition from network-level processing to local processing and ultimately hindering the transformation and consolidation of emotional memories during sleep.

### 1.3. The current study

This study aimed to determine whether intrinsic aperiodic slopes influence sleep-related emotional memory consolidation from childhood to late adulthood. Understanding this relationship will likely advance our knowledge of the functional maturation of the human brain and how sleep and emotional processing may influence normative functioning across the lifespan. Furthermore, the phenomenon of emotional enhancement, while beneficial to long-term memory processes, is considered a significant factor in the etiology of several mental health conditions, including depression, anxiety, and post-traumatic stress disorder (Philippot, Schaefer & Herbette, 2003). In this context, the vividness of memories with greater emotional tone may be paramount to better approximating that particular range of experiences. It is also important to consider that a sleep-related bias in emotional information processing may contribute to these forms of psychopathology (Everaert et al., 2022).

We employed a repeated-measures, within-subjects design, where participants aged 7–72 years of age completed both sleep and wake conditions to control for individual differences and condition-specific effects. Further, rather than relying on arbitrary valence ratings, we used ratings of stimuli from participants, strengthening the accuracy and relevance of emotional categorization by accounting for individual differences in emotional perception. We also used advanced spectral decomposition techniques (Wen & Liu, 2016) to separate aperiodic from oscillatory activity, alongside analyses of sleep EEG microstructure (Vallat & Walker, 2021), including spindle and SO activity, markers of sleep-based memory consolidation (Denis & Cairney, 2024) that differ with age (Mander, Winer, & Walker, 2017).

Based on published work on age-related differences in sleep and memory consolidation (Gui et al., 2017) and aperiodic neural activity (Cross et al., 2024; Merkin et al., 2023; Thuwal et al., 2021), we predicted that: (1) age-related differences in the aperiodic slope will predict the consolidation of emotionally valenced relative to neutral information, and this effect will be stronger after sleep relative to wake; (2) this effect will be strongest for negatively valenced information in children and young adults, and positively valenced information in older adults. We also performed exploratory analyses to establish whether the interaction between resting-state slopes and sleep microstructural activity predicts age-related differences in emotional memory.

## 2. Methods

### 2.1. Participants

Participants included 42 healthy individuals ranging from 7–72 years old (*M* = 26.60, *SD* = 17.45; 26 female). All participants reported normal or corrected-to-normal vision and hearing, were native English speakers, and were predominantly right-handed. No participants reported ongoing medication, health problems or a history of psychiatric, neurological, or sleep conditions. No adult participants had a history of substance abuse or recreational drug-use within the past 6 months. Furthermore, all participants were rated as good sleepers in the month prior to the study as rated by the Pittsburgh Sleep Quality Index (*M* = 3.00, *SD* = 1.49) for adults, or the Sleep Disturbance Scale for children (*M* = 39.71, *SD* = 4.38). All participants (and their legal guardians in the case of children) provided informed consent and received an $80 honorarium. Ethics for this study was granted by the University of South Australia’s Human Research Ethics committee (I.D: 0000032556, 0000034184).

### 2.2. Study Design

This study adopted an experimental, repeated measures, within-subjects design with two conditions (sleep, wake). The order of condition was randomized to control for order effects and separated by a week to avoid interference. All participants were required to complete both conditions. Conditions included (see Figure 1):

**Figure 1.**
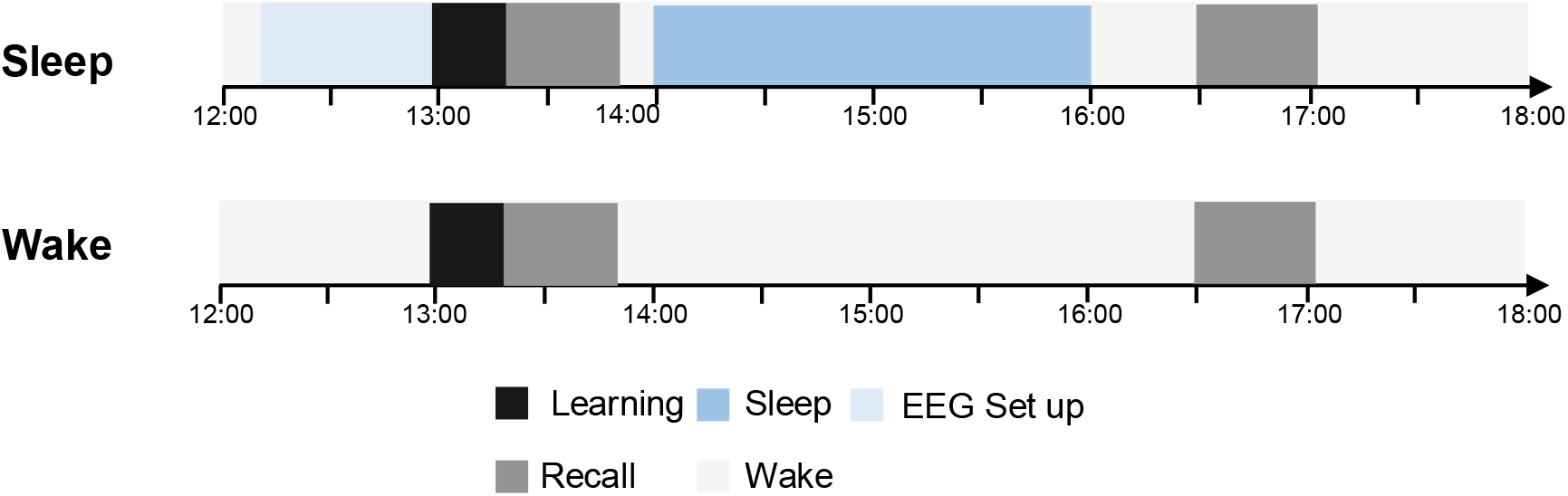
Diagram representing the time course (24hr clock time) of the experimental conditions (Sleep, Wake) and testing session (Learning, Recall).

a. Sleep (nap) condition: participants underwent learning with a baseline recall task followed by a 2hr afternoon sleep opportunity. A delayed recall task occurred thirty minutes after waking.
b. Wake condition: participants underwent learning with a baseline recall task. This was followed by a delayed recall task after a 2hr wake period.

### 2.3. Demographic and control measures

Participants completed a questionnaire containing questions on age, sex, ethnicity, highest level of education achieved, and for adults, recent (<24hr) alcohol and caffeine consumption.

#### 2.3.1. Handedness

The Flinders Handedness Scale (FLANDERS) was used as a screening measure of handedness, as hand preference is known to be an indicator of functional cerebral lateralization (Nicholls, Thomas, Loetscher, & Grimshaw, 2013). The FLANDERS demonstrates very high split-half reliability for hand preference (Cronbach’s alpha = .96; Nicholls, Thomas, Loetscher, & Grimshaw, 2013).

#### 2.3.2. Sleep quality

Participants completed a measure of sleep disturbance and regular sleep habits. Adult participants completed the Pittsburgh Sleep Quality Index (PSQI). The PSQI contains 19 self-rated questions, making up seven component scores: subjective sleep quality, sleep latency, sleep duration, habitual sleep efficiency, sleep disturbances, sleeping medication, and daytime dysfunction (Buysse et al., 1989). Scores are added to give a global score out of 21, where <5 is associated with good sleep quality and >5 is indicative of poor sleep quality. Those with a score higher than 5 were excluded from the study. The PSQI is reported to have high internal consistency and has a reliability coefficient of α=.83 for its components.

Parents of child participants completed the Sleep Disturbance Scale for Children (SDSC). The SDSC is a 26-item parent-report measure used to examine six domains of sleep behavior and problems in children aged 6-15 years: disorders of initiating and maintaining sleep, sleep breathing disorders, disorders of arousal/nightmares, sleep-wake transition disorders, disorders of excessive somnolence, and sleep hyperhidrosis (i.e. night sweats; Bruni et al, 1996). The parent/caregiver of the participant was asked to indicate the typical frequency in which their child has engaged in certain behaviours. Global scores of < 51 were excluded from the study, as it is indicative of a clinical sample with high difficulties in sleep behavior and sleep quality. The SDSC has demonstrated a diagnostic accuracy of α=.91, test-retest reliability of α=.71, and reliability coefficient ranging from α=.71 to α.=79 in control and clinical samples, respectively (Bruni et al, 1996).

#### 2.3.3. Intelligence

The Wechsler Abbreviated Scale of Intelligence - 2nd edition (WASI-II) was used to control overall cognitive ability, as intelligence may influence memory retention and performance on memory tasks (Alexander & Smales, 1997; Conway, Kane & Engle, 2003). The WASI-II is a standardized battery of four subtests used to gain an estimate of full-scale IQ (FSIQ-4; Wechsler, 2011). The WASI-II has normative samples for ages 6 to 90 years, with a reliability score of (α=.97) for FSIQ-4, and concurrent validity established with alternate measures of intelligence such as the Wechsler Adult Intelligence Scale - Fourth Edition and the Wechsler Intelligence Scale for children - Fourth Edition (Wechsler, 2011). Participants’ mean FSQI score was 104.14 (*SD* = 9.24), placing participants in an average to high range of intellectual functioning (Wechsler, 2011).

#### 2.3.4. Sleepiness

Measures of sleepiness were in the form of a 100mm visual analogue scale with “sleepy/drowsy” and “alert/awake” for endpoints to denote a continuum of state sleepiness. Participants indicated on the line where they judge their current state of sleepiness to be at time of testing. Participants were asked to complete the visual analogue scale for sleepiness (VASS) prior to the start of each encoding and testing session. Degree of sleepiness is considered as the distance from the left pole to the participants’ mark, indicating a percentage. VASS measures of sleepiness are typically used in research settings where temporal resolution of state changes are being examined, thus suitable for looking at within-subject changes across time (Drake, 2011).

#### 2.3.5. Valence and arousal of stimuli

The Self-Assessment Manikin (SAM) is a pictorial assessment technique used to measure an individual’s affective reactions to emotionally valenced stimuli (Bradley et al., 1992; Bradley & Lang, 1994). It is a widely used, inexpensive and easy method for assessing affective responses in both adults and children (Bradley & Lang, 1999; Lang, Bradley & Cuthbert, 2008; McManis, Bradley, Berg, Cuthbert, & Lang, 2001). Participants completed a paper questionnaire containing all stimuli presented in each condition. The questionnaire included two rating scales per stimulus on a 9-point scale, representing emotional valence, i.e. 9 (happy) to 1 (unhappy), and subjective arousal, i.e. 9 (excited) to 1 (relaxed). On presentation of each stimulus, participants were required to indicate on each scale where their immediate experience fell while viewing the stimulus.

### 2.4. Emotional memory task

Affective stimuli were adapted from Bennion and colleagues (2014). The stimuli are static visual images composed of valenced foreground objects, plausible neutral backgrounds, and composite scenes of the background and foreground objects. The term ‘plausible’ here denotes that the foreground objects could theoretically be observed on the backgrounds in real life. All stimuli were previously rated on valence. However, participants in the current study rated their own levels of valence and arousal on a self-assessment manikin (SAM; Bradley & Lang, 1999) post-condition. Affective stimuli were presented using OpenSesame Software version 3.0.5 (Mathôt, Schreij & Theeuwes, 2012). 240 composite images were used, which were divided into two parallel sets, counterbalanced across the sleep and wake conditions. Both children and adult participants viewed the same test stimuli.

During the learning tasks (see Figure 3A for schematic diagram), 90 composite images were presented. Composite images consisted of a valenced (positive, negative, neutral) foreground image, on a neutral background. Stimuli were presented for 1000ms to allow conscious processing, while maintaining an appropriate degree of difficulty in order to prevent ceiling effects at later testing times. After this presentation time participants were required to indicate whether they would approach or back away from the scene if they encountered it during real life. Stimuli were preceded by a 500ms fixation cross and had an interstimulus interval (ISI) of 1500ms.

Following the learning task there were two recall sessions (see Figure 3A), each with 360 stimuli (180 backgrounds, 180 foreground objects; 90 targets/90 distractors). The first recall session was carried out immediately, in order to achieve a recognition baseline to control for level of encoding. The delayed recall session took place after the retention period of sleep or wake. In all retrieval sessions, participants were presented with foreground objects and backgrounds separately. Each stimulus was presented until a response was given via a button press. After the presentation time, participants indicated whether the stimulus was old (previously presented in encoding) or new (a distractor item not seen in the learning context of the experiment). Next, participants indicated recognition confidence on a 10-point Likert scale, ranging from 0 (not confident) to 10 (absolutely confident), as seen in Weymar and colleagues (2009; 2011). Stimuli were pseudo-randomized at each testing time so that they were intermixed such that no more than two stimuli of the same content or emotion followed each other.

### 2.5. Procedure

Consenting participants came to the laboratory at approximately 11:45hr and were taken through to the testing rooms for EEG set-up. Prior to the learning task participants completed the VASS. Resting-state EEG was recorded during quiet sitting with eyes open (focusing on a fixation cross centered on a computer monitor) and eyes closed for two minutes, respectively (See Figure 3B for schematic of resting-state and sleep EEG). Participants then received the learning tasks at approximately 13:00hr, followed by immediate recall tasks (baseline). During the sleep condition, participants were given a 120-minute sleep opportunity between the hours of approximately 14:00hr and 16.00hr. During the wake condition, participants then remained in the laboratory and were administered the WASI-II. PSG data were used to verify sleep periods. Participants engaged in non-strenuous activity for 30 minutes after the nap until testing to alleviate inertia effects on memory performance. At approximately 16:30hr, participants completed the VASS again and then given the delayed recall task (see Figure 1 for study protocol). Post testing participants were given the paper SAM questionnaire for stimuli presented in that condition.

## 3. Data analysis

### 3.1. Behavioral data

D prime (d’) was used as an indicator of overall memory performance in accordance with Signal Detection Theory (SDT; Macmillan & Creelman, 1991; Stanislaw & Todorov, 1999). D’ represents a sensitivity index that reflects the distance between the signal and signal+noise distributions (i.e., how well a participant can distinguish targets from distractors) and is calculated as the difference between standardized hit rate (HR) and false alarms (FA; incorrectly identified distractors, i.e., zHR-zFA). Adjustment of extreme values was made using the recommendations of Hautus (1995). These scores were calculated for each retrieval session (baseline, delayed), in addition to across conditions (sleep, wake) and valence categories (positive, negative, neutral).

### 3.2. Polysomnography (PSG)

Continuous EEG activity was recorded during nap opportunities using a BrainAmp DC system (BrainAmp MR Plus, Brain Products) with a 32-channel BrainCap with sintered Ag/AgCI electrodes. Additional electrodes included bipolar horizontal electrooculogram (EOG:LOC/ROC placed on the left and right outer canthi, respectively), and submental electromyogram (EMG), for standard monitoring of sleep periods. EOG, EMG, and EEG were recorded using a band-pass filter of 0.1–100Hz and sampled at a rate of 1000Hz. All impedances were maintained at or kept below 15kΩ throughout recording periods.

All sleep data were scored by an experienced sleep technician to the AASM criteria described by Berry and colleagues (2012). EEG data were re-referenced offline to contralateral mastoids. The following sleep parameters were derived from PSG recordings: total sleep time (TST), sleep onset latency (SOL; time from lights out to the first epoch of sleep), wake after sleep onset (WASO), minutes and percent of TST spent in each sleep stage (N1, N2, N3, and REM).

### 3.3. Resting-state aperiodic slope estimation

To extract the aperiodic slope from resting-state EEG recordings, we used the irregular-resampling auto-spectral analysis method (IRASA; Wen & Liu, 2016) implemented in the YASA toolbox (Vallat & Walker, 2021). IRASA isolates the aperiodic component of neural time series data via a process that involves resampling the signal at multiple non-integer factors *h* and their reciprocals 1/*h*. This resampling procedure systematically shifts narrowband peaks away from their original location along the frequency spectrum. Averaging the spectral densities of the resampled series attenuates peak components, while preserving the 1/*f* distribution. The exponent summarizing the slope of aperiodic spectral activity is then calculated by fitting a linear regression in log–log space. See Figure 2 and Figure 3C (top) for a schematic of the separation of aperiodic and oscillatory activity and varying slope estimates, respectively.

**Figure 2.**
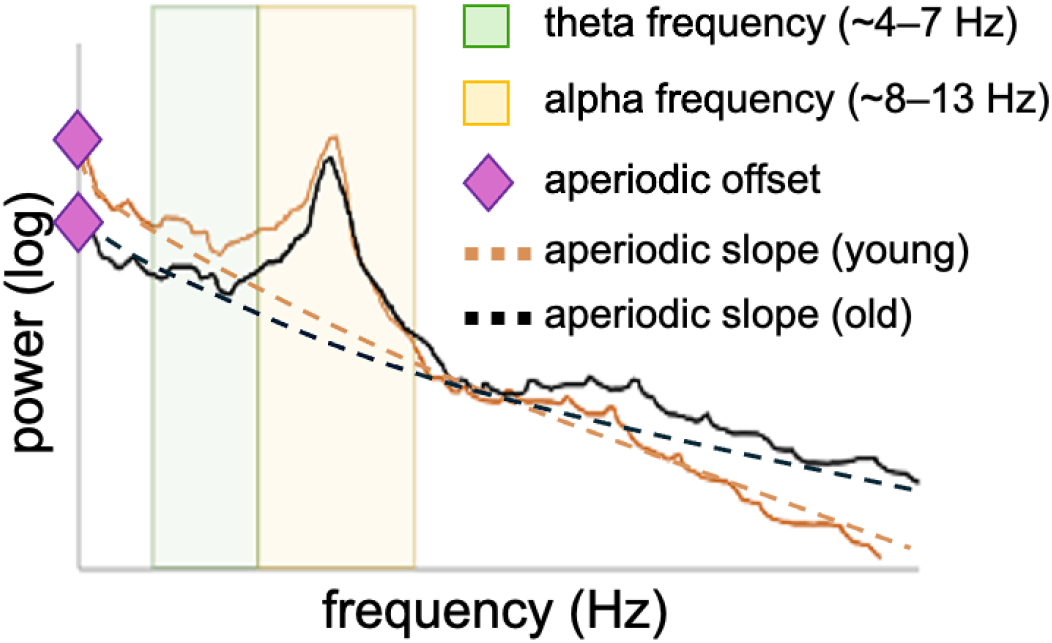
Schematic of the separation of aperiodic (curved dashed lines) and periodic (oscillatory; solid lines) activity in power spectral density space. Older individuals typically exhibit a flatter slope (dashed black line) and lower offset, while younger individuals typically have a steeper slope (dashed orange line) and a higher offset. The canonical theta and alpha frequencies are shown in green and yellow shaded regions, with alpha showing a prominent peak at approximately 10 Hz.

### 3.4. Sleep microstructural EEG estimation

In addition, spindle density, SO density, and SO-sigma phase-dependent correlations were extracted using the YASA (*Yet Another Spindle Algorithm*) toolbox (Vallat & Walker, 2021) based on published algorithms (Helfrich et al., 2018; Mölle, Bergmann, Marshall, & Born, 2011; Staresina et al., 2015; see Figure 3C bottom). The EEG data were re-referenced to linked mastoids and a true-finite impulse response bandpass filter was applied (0.3–30 Hz). Artifact rejection was conducted on sleep staged data based on covariance-based rejection methods with the hypnograms (Vallat & Jajcay, 2020). Epochs were rejected based on a kurtosis threshold >3.

**Figure 3.**
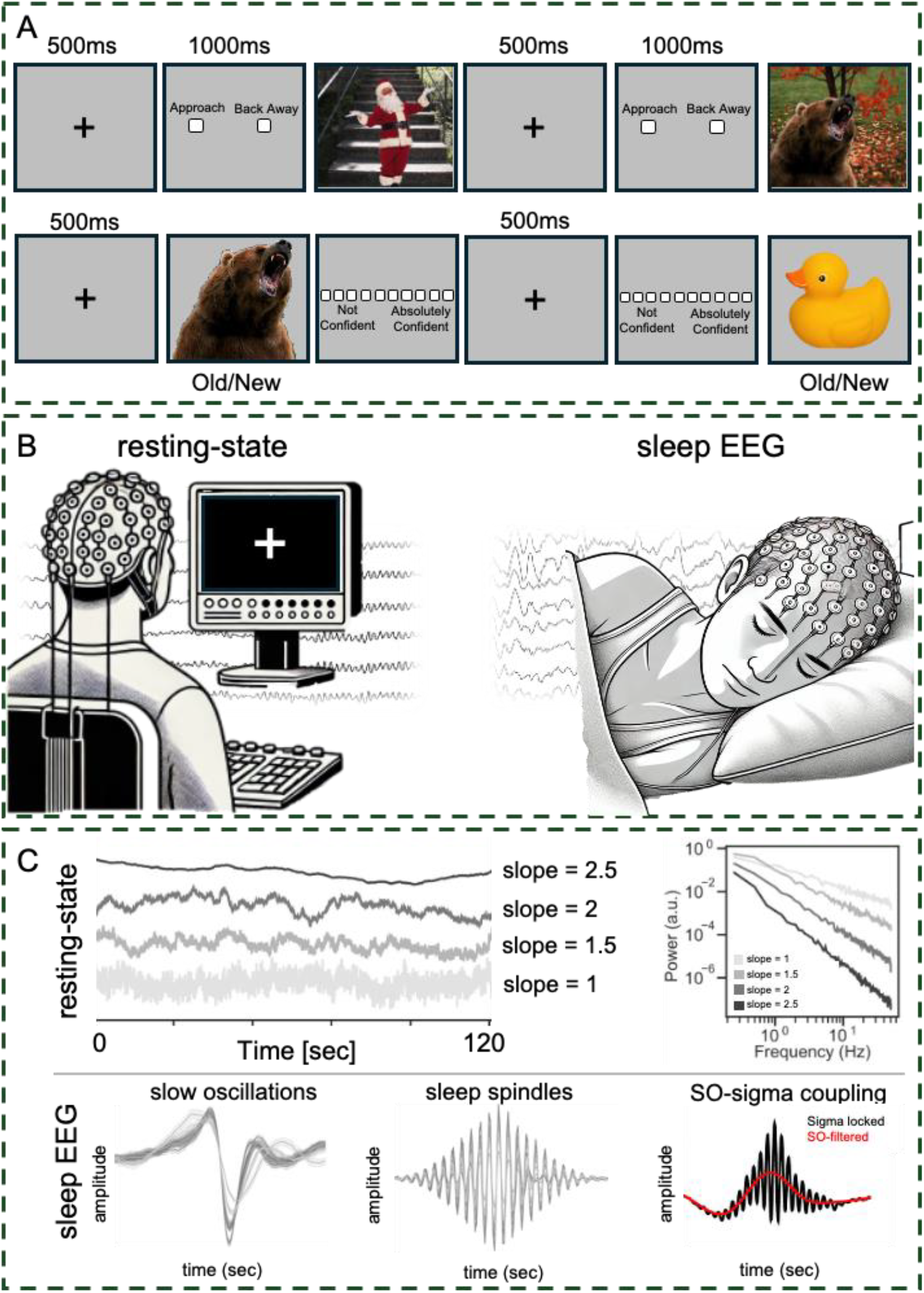
Schematic of the emotional memory task, EEG recording paradigm, and resting-state and sleep EEG microstructural activity. **(A)** Illustration of the visual memory task adapted from Bennion et al (2014). Top; schematic representation of the learning task. Each stimulus is a composite image with a valenced (positive, negative, neutral) foreground image, on a neutral background. Stimuli were presented for 1000ms, after which they were required to indicate whether they would approach or back away from the scene if they encountered it during real life. The interstimulus interval was 1500ms. Bottom: Schematic representation of the recall task. Participants were presented with foreground objects and backgrounds separately during recall. Each stimulus was presented until button press. During the presentation time, participants needed to indicate whether the stimulus was old or new. Participants were also required to provide an indication of confidence for each response. **(B)** Illustration of the resting-state and sleep-EEG recording sessions. **(C)** Top left: depiction of ongoing EEG activity across a two minute (120 second) recording with corresponding aperiodic slope values. Top right: decomposition of the aperiodic slope values into a power spectral density plot (adapted from Schmidt et al., 2024). Bottom left: schematic illustration of the main microstructural sleep EEG variables: slow oscillations, sleep spindles, and slow oscillation-sigma coupling.

Spindle detection algorithms were conducted with the YASA spindle algorithm based on Lacourse et al. (2019) and Purcell et al. (2017). Sigma frequency range (11–16 Hz) relative to the total power band frequency (1–30 Hz) was calculated using a short-term Fourier transform on consecutive epochs of 2s and with an overlap of 200ms. The root mean square was then computed with a window size of 300ms and a step of 100ms. The resulting signal was then smoothed in the same window with a moving average. Amplitude threshold was set at 75% (amplitude criterion) and events were only included if they were longer than 0.5–3s (time criterion). Spindle density (i.e., *n* spindles per 30s NREM epoch) was then calculated for each participant at channel Cz.

Slow oscillation detection algorithms were conducted with YASA based on Massimini et al. (2004) and Carrier et al. (2011). For SOs, continuous EEG data were filtered from 0.16–1.25Hz to detect zero crossing events that were between 0.8–2s in length, and that met a 75-microvolt criterion. These artifact-free epochs were then extracted in segments of 5s (± 2.5s of SO trough) from the raw EEG signal. Artifact-free events were then defined as 5s (± 2.5s) peak-locked sigma power epochs. An event-locked cross-frequency coupling metric was calculated (for a detailed description of this method, see Helfrich et al., 2018). For every participant at channel Cz and epoch, maximal sigma amplitude and corresponding SO phase angle was detected. The mean circular direction (coupling phase) and resultant vector length (coupling strength) across all NREM events were determined using the Pingouin statistical package in Python (Vallat, 2018). Coupling phase represents the SO phase in degrees (i.e. with 0° being at the positive peak of the SO) where the sigma amplitude is at its maximal. Coupling strength is represented by normalized direct phase-amplitude cross-frequency coupling (ndPAC). This is assessed using a scale of 0–1, where 0 indicates every coupled sigma envelope occurred towards a different phase of the SO, as opposed to 1 which indicates coupled sigma activity occurred in the same SO phase.

### 3.5. Statistical analysis

Data were imported into *R* version 4.2.3 (R Core Team, 2020) and hypotheses were tested using linear mixed-effects models fit by restricted maximum likelihood (REML) using *lme4* (Bates, 2010). To examine age-related differences in recognition accuracy as a function of regional (i.e., across the scalp) aperiodic activity, valence, and condition, we constructed the following model:

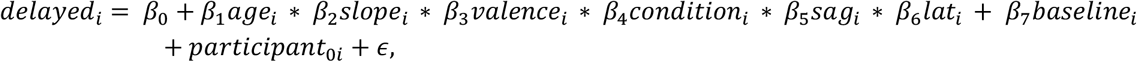

with *delayed* being recognition accuracy (d’ prime) post-sleep or wake, *age* is age in years, *slope* is the resting-state aperiodic slope values, *valence* encodes the emotional valence (negative, positive, neutral), *condition* refers to the sleep or wake conditions, *sag* encodes the sagitality of the aperiodic slope estimates (anterior, central, posterior), while *lat* refers to the laterality of the slope estimates (left, midline, right); *baseline* is recognition accuracy prior to the sleep or wake period. *Participant* is participant modelled as a random effect on the intercept, while *ε* indicates a Gaussian error term.

For sleep-related analyses, we constructed separate linear mixed-effects models to predict recognition accuracy for each sleep EEG predictor, with sleep EEG (i.e., spindle density, SO density, SO-sigma coupling strength, or SO-sigma coupling phase), age (in years), aperiodic slopes, and valence (negative, neutral, position), with full interactions between these variables. We also covaried for baseline recognition accuracy (Alday, 2019) and specified participant as a random effect on the intercept.

Effects were plotted using the package *effects* (Fox & Hong, 2010) and *ggplot2* (Wickham & Wickham, 2016). Post hoc comparisons for main effects were performed using the *emmeans* package (Lenth et al., 2023). Categorical factors were sum-to-zero contrast coded, such that factor level estimates were compared with the grand-mean (Schad et al., 2020). Further, for modeled effects, an 83% confidence interval (CI) threshold was used given that this approach corresponds to the 5% significance level with nonoverlapping estimates (Austin and Hux, 2002; MacGregor-Fors and Payton, 2013). In the visualization of effects, non-overlapping CIs indicate a significant difference at *p* < 0.05.

## 4. Results

### 4.2. Preliminary analyses

Preliminary analyses were conducted to determine: (1) if there was a difference in subjective sleepiness between the conditions prior to baseline learning and delayed recall tasks; (2) the sleep characteristics of the nap; (3) if there were differences in valence and arousal ratings between the normative values and participants’ self-ratings; (4) the aperiodic slope and sleep EEG characteristics, and (5) recognition accuracy characteristics between conditions and valences.

#### 4.1.1. Subjective sleepiness

No difference was found for subjective sleepiness prior to baseline testing for the sleep (*M* = 56.95, *SD* = 19.49) and wake (*M* = 57.22, SD = 21.75) conditions (*t*(39) = -.16, *p* = .87, *d* = -.01). There was also no difference in subjective sleepiness between conditions prior to delayed recall (sleep: *M* = 64.12, *SD* = 23.03, wake: *M* = 60.34, *SD* = 24.28; *t*(40) = 0.95, *p* = .35, *d* = .16).

#### 4.1.2. Nap characteristics

A summary of sleep parameters including TST, SOL, WASO and the amount of time and percentage of sleep spent in stage 1 (N1), stage 2 (N2), SWS and REM are reported in Table 1. Sleep data for young and middle adults show comparable patterns of NREM sleep stages (i.e., N1, N2 and SWS) and marginally greater amounts of REM sleep to that of Payne and colleagues (2015) and Alger and colleagues (2018). However, the current study used a 120-minute nap opportunity, as opposed to 90 minutes in each of those studies (Alger et al., 2018; Payne et al., 2015). Nap success rate was 95% in the current study, although only 52% of participants achieved REM sleep. Further specified by age range, 43% of those aged 7–17 years achieved REM, compared to 67% of those aged 18–45 and 29% of those aged 46–72 years.

**Table 1.** Means, standard deviations, and ranges of sleep parameters for all ages.

| Parameter | Mean | Standard Deviation | Range |
| --- | --- | --- | --- |
| TST (min) | 70.36 | 34.48 | 0.00 - 116.00 |
| SOL (min) | 14.83 | 20.39 | 1.00 - 119.00 |
| SE (%) | 65.65 | 24.78 | 10.20 - 95.90 |
| WASO (min) | 21.46 | 22.54 | 1.00 - 82.50 |
| N1 (min) | 6.24 | 5.71 | 0.00 - 29.50 |
| N1 (%TST) | 8.34 | 7.33 | 0.00 - 40.00 |
| N2 (min) | 35.85 | 20.89 | 0.50 - 70.00 |
| N2 (%TST) | 48.52 | 19.42 | 0.80 - 87.80 |
| N3 (min) | 24.75 | 19.00 | 0.00 - 76.50 |
| N3 (%TST) | 35.16 | 25.82 | 0.00 - 95.80 |
| REM (min) | 6.93 | 9.94 | 0.00 - 38.00 |
| REM (%TST) | 7.15 | 10.18 | 0.00 - 36.60 |

*Note.* TST = total sleep time; SOL = sleep onset latency; WASO = wake after sleep onset; N1 = stage 1; N2 = stage 2; N3 = slow wave sleep; REM = rapid eye movement sleep; min = minutes.

#### 4.1.3. Ratings of valence and arousal

Cohen’s Kappa was calculated to determine if there was agreement between classification of stimuli in each valence category (negative, neutral, positive) between the current participants and established categories from Bennion et al. (2014). There was excellent agreement beyond chance for valence judgements (κ = .88, 95% CI, .84 to .92, *p* <.001), with the percentage of agreement found to be 92.2%. The following analyses use valence categories based on self-rated scores.

#### 4.1.4. EEG characteristics

See Table 2 for a summary of the ranges and bivariate correlations with age and recognition accuracy (collapsed across valence categories) for the key sleep and resting-state aperiodic EEG variables. For SO-sigma coupling, there was a significant nonuniform distribution for the precise SO phase during peak sigma activity (*p*≤0.001; Rayleigh test).

**Table 2.** Ranges and bivariate correlations between sleep EEG and aperiodic variables with age and memory.

| EEG Variables | Range | Correlation with Age<br>$r$ ($p$) | Correlation with Memory<br>$r$ ($p$) |
| --- | --- | --- | --- |
| **Sleep Microstructural EEG** |  |  |  |
| SO Density | 0.34 – 15.09 | **-0.70 (<0.001)** | -0.01 (0.95) |
| Spindle Density | 0.02 – 2.41 | 0.15 (0.41) | 0.28 (0.13) |
| Coupling Strength | 0.13 – 0.24 | 0.27 (0.14) | 0.06 (0.74) |
| Coupling Phase | -1.73 – 2.26 | -0.23 (0.22) | 0.09 (0.61) |
| **Resting Aperiodic EEG** |  |  |  |
| Slope | 1.87 – 3.03 | **-0.63 (<0.001)** | 0.18 (0.28) |
| Offset | -24.31 – -18.97 | **-0.76 (<0.001)** | -0.20 (0.24) |
| Model Fit ($R^2$) | 0.47 – 0.77 | **-0.52 (<0.001)** | -0.07 (0.70) |

*Note.* Correlation with memory represents correlations with recognition accuracy collapsed across valence categories; offset = the intercept of the aperiodic model fit, reflecting the overall power law; model fit ( $R^2$ ) indicates the goodness-of-fit of the aperiodic model. Significant correlations are boldfaced. Correlations are spearman rho coefficients; $r$ = correlation coefficient; $p$ = statistical significance.

#### 4.1.5. Recognition accuracy differences

The results of the baseline and delayed recall sessions are given in Table 3. The sleep condition had a higher overall d’ score at delayed testing compared to the wake condition, particularly for negatively and positively valenced stimuli.

**Table 3.** Mean d’ scores by condition (sleep, wake), time (baseline, delayed) and valence (positive, negative, neutral). Standard deviations are given in parentheses.

| Condition | Time | Valence | Mean (SD) |
| --- | --- | --- | --- |
| Sleep | Baseline | Negative | 2.04 (1.01) |
|  |  | Neutral | 1.68 (0.87) |
|  |  | Positive | 2.08 (1.09) |
|  | Delayed | Negative | 1.68 (1.03) |
|  |  | Neutral | 1.47 (0.89) |
|  |  | Positive | 1.79 (1.01) |
| Wake | Baseline | Negative | 1.85 (1.08) |
|  |  | Neutral | 1.60 (0.96) |
|  |  | Positive | 1.95 (1.09) |
|  | Delayed | Negative | 1.67 (0.92) |
|  |  | Neutral | 1.43 (0.99) |
|  |  | Positive | 1.57 (0.89) |

### 4.2. Recognition accuracy and aperiodic activity differ across the lifespan

In a first step, we sought to establish age-related differences in emotional memory and intrinsic (resting-state) aperiodic activity. For the analysis of emotional memory, we implemented a linear regression with a polynomial spline (two internal knots) on Age. In doing so, we revealed a nonlinear main effect of Age, such that recognition accuracy increased from approximately seven to 30 years of age, decreasing thereafter until 72 years of age (*β* = -0.80, *SE* = 0.29, *p* = .008; Figure 4A). This analysis did not reveal a main effect of Valence (*β* = -0.10, *SE* = 0.26, *p* = .71) or an Age × Valence Interaction (*β* = 0.54, *SE* = 0.69, *p* = .43) on recognition accuracy. While no differences in memory between emotional valence categories may be somewhat surprising given prior literature (Kensinger, 2009), it is important that we consider interindividual differences in information processing capacities across the lifespan, such as interindividual differences in aperiodic neural activity. We address this in Sections 4.3. and 4.4. after establishing lifespan variability in aperiodic slopes, as well as condition (sleep, wake) differences in emotional memory.

**Figure 4.**
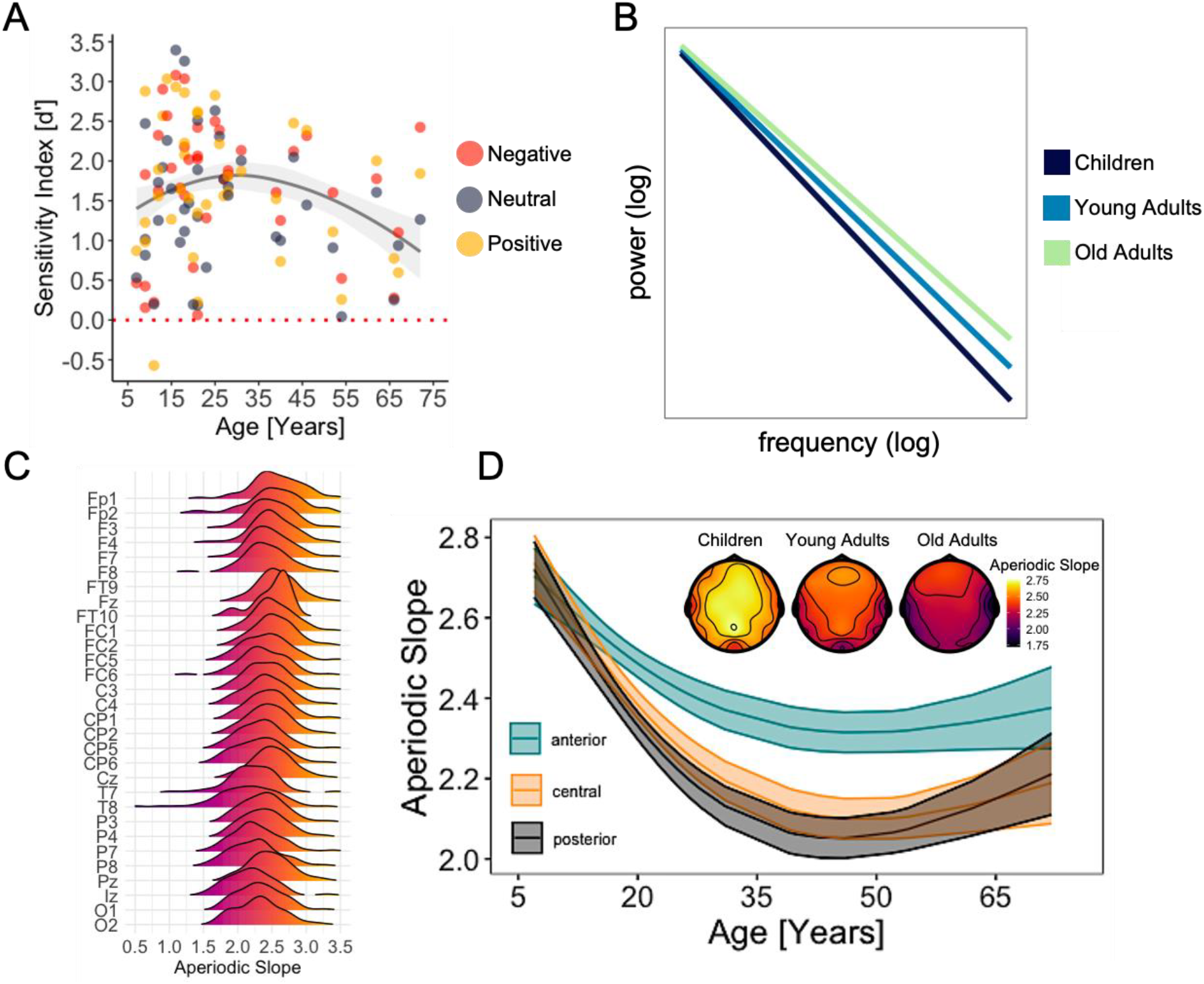
**(A)** Modelled relationship between memory (d’; y-axis, higher values denote better memory) and age (x-axis). Individual data points represent single participant memory performance values for negative (red), neutral (gray) and positive (yellow) stimuli. The dashed red line indicates chance performance, while the shaded region indicates the 83% confidence interval. **(B)** Illustration of aperiodic slopes (negative exponent) in log-log space for children (black), young adults (blue) and older adults (green). **(C)** Ridgeline plot illustrating the distribution of aperiodic slopes (x-axis; higher values denote a steeper slope) by channel (y-axis). **(D)** Modelled effects for differences in the aperiodic slope (y-axis; higher values denote a steeper slope) and age (x-axis) across anterior (teal), central (orange) and posterior (black) channels. Shaded regions indicate the 83% confidence interval. Topoplots are presented to show the global distribution of aperiodic slopes across age (warmer colors denote a steeper slope).

For aperiodic activity, we modelled the aperiodic Slope as a function of Age (again, with a polynomial spline), Sagitality (anterior, central, posterior) and Laterality (left, midline, right), revealing a nonlinear main effect of Age (*β* = -0.99, *SE* = 0.08, *p* < .001; see Figure 4C for distributions of slopes across channels) and an Age × Sagitality interaction (*β* = 0.31, *SE* = 0.11, *p* = .005). As is clear from Figure 4D, aperiodic slopes are steepest in childhood, flattening until approximately 50 years of age, before steepening thereafter; this effect is most prominent in anterior regions. This analysis indicates that aperiodic slopes demonstrate a u-shaped curve across the lifespan and are steepest in anterior regions, consistent with recent intracranial EEG work (Cross et al., 2024).

### 4.3. Age-related differences in emotional memory are modulated by sleep-wake differences

Having established that memory and aperiodic slopes differ across the lifespan, our next step was to examine whether age-related differences in emotional memory are differentially influenced by sleep versus wake periods and interindividual differences in aperiodic slopes. To achieve this, we constructed a linear mixed-effects regression with delayed Recognition Accuracy as the outcome, and Slope, Age, Valence (positive, negative, neutral), Condition (sleep, wake), Sagitality (anterior, central, anterior), and Laterality (left, midline, right) as fixed effects. Baseline Recognition Accuracy was specified as a covariate (Alday, 2019), and Participant as a random effect on the intercept.

Here, we focus on subordinate main effects and low-level interactions. This model revealed a significant main effect of Valence, whereby memory for negative stimuli was better than for neutral and positive stimuli (χ2(2) = 14.80, *p* < .001; Figure 5A). There was also an Age × Valence interaction, such that memory decreased for neutral and positive stimuli as age increased, whereas no age-related decreases in memory were observed for negative stimuli (χ2(2) = 20.74, *p* <.001; Figure 5B). Finally, the model revealed a significant Condition × Valence interaction (χ2(2) = 49.30, *p* <.001; Figure 5C), whereby memory was overall higher for negative stimuli across conditions, while memory for positive stimuli was higher in the sleep relative to the wake condition. While these results differ from those reported in Section 4.2., the inclusion of Condition (sleep, wake) and Slope (a marker of interindividual, age-related differences in information processing capacities) allows for a better partitioning of variance. This approach accounts for both the consolidation period and individual differences in neural processing, offering a more nuanced understanding of how these factors interact to influence age-related differences in emotional memory.

**Figure 5.**
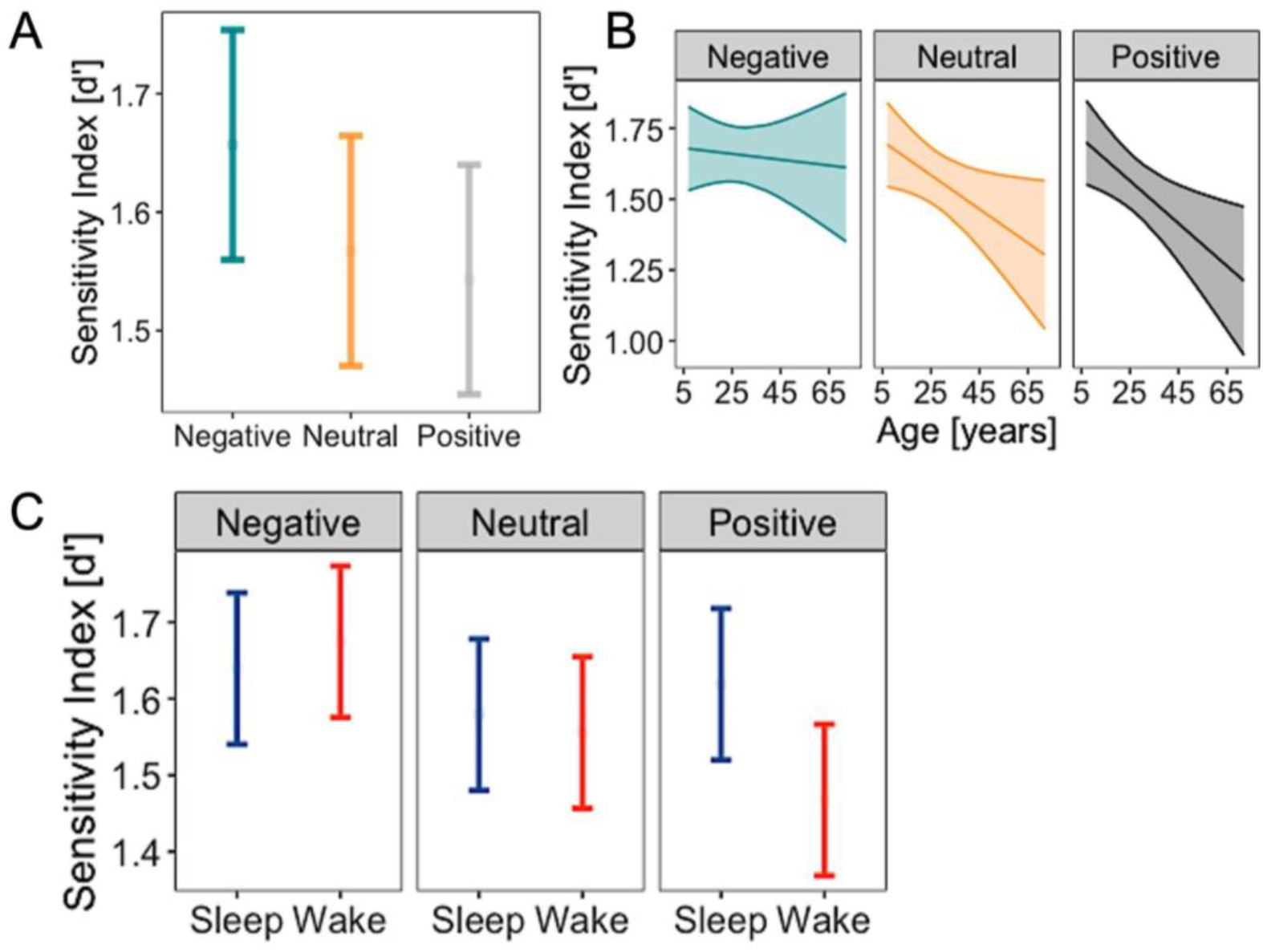
**(A)** Modelled relationship between d’ scores at delayed testing (y-axis; higher scores denote better memory) and valence (x-axis; negative, neutral, positive). **(B)** Estimated marginal means for d’ scores at delayed testing by age (x-axis) and valence (negative = left, neutral = middle, positive = right). Shaded regions indicate the 83% CI. **(C)** Modelled relationship between d’ scores at delayed testing (y-axis; higher scores denote better memory), condition (x-axis; sleep = blue, left; wake = red, right) and valence (negative = left, neutral = middle, positive = right).

### 4.4. Age-related flattening of the aperiodic slope weakens sleep-based emotional memory consolidation

Here, we focus on the highest-level interaction in the model described above in Section 4.3., which was revealed to be an Age × Slope × Valence × Condition interaction (χ2(2) = 21.00, *p* < .001; Figure 6). As shown in Figure 5, in the top-left panel representing the sleep condition and negative stimuli, a clear divergence between steep (orange; less neural noise) and flat (teal; more neural noise) slopes is observed. Steeper slopes exhibit a positive relationship between age and memory, with memory improving as age increases. This suggests that older individuals with steeper slopes have superior memory for negative stimuli after sleep. In contrast, flatter slopes show a negative relationship between age and memory, with memory declining as age increases, indicating that individuals with flatter slopes experience a more substantial decline in memory for negative stimuli with age. This effect highlights that the relationship between age and memory for negative stimuli is modulated by resting-state aperiodic slopes, with steeper slopes associated with a more pronounced retention of negatively toned memory across the lifespan. Less pronounced effects are observed across the remaining valence and condition levels as a function of age.

**Figure 6.**
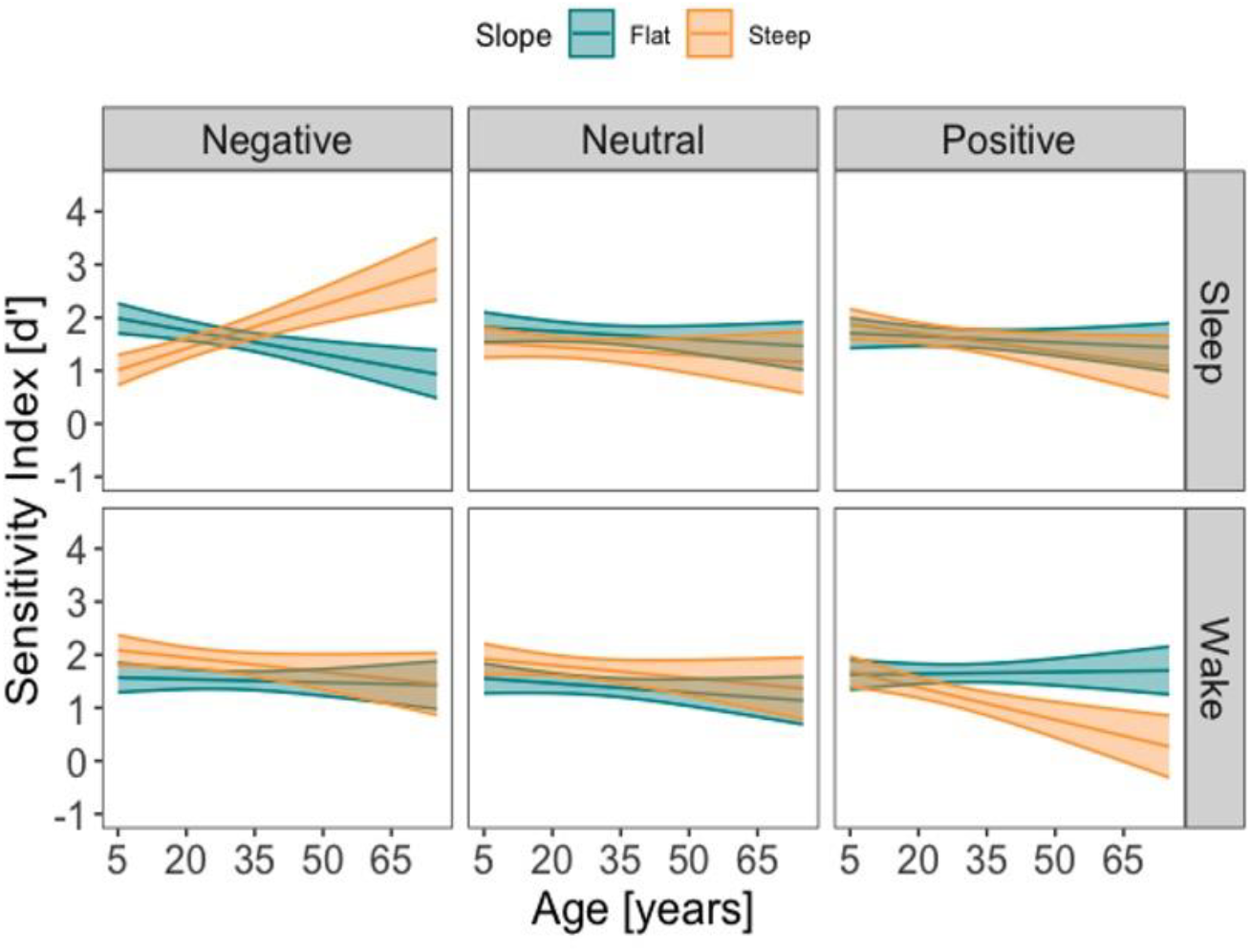
Modelled effects of age (x-axis; in years) on d’ scores at delayed testing (y-axis; higher values indicate better memory) as a function of valence (negative = left, neutral = middle, positive = right) and condition (sleep = top row; wake = bottom row). Resting-state aperiodic slopes are dichotomized into flat (teal) and steep (orange) for visualization purposes only, with the aperiodic slope being entered into all models as a continuous predictor. The shaded regions indicate the 83% CI.

### 4.5. NREM slow oscillatory activity predicts age-related differences in emotional memory

Having demonstrated that age-related differences in aperiodic slopes modulate memory for negative information in the sleep condition, we next aimed to determine whether slopes similarly interact with sleep microstructural EEG to influence emotional memory outcomes. For clarity, we only report significant main effects and/or interactions that involve sleep EEG and aperiodic slopes. While there were no such effects observed in the models examining NREM sleep spindle density and SO-sigma coupling strength, we observed a significant SO Density × Valence interaction (χ2(2) = 10.70, *p* = .005; Figure 7A). Here, as SO density increased (i.e., more SO events per 30s of NREM sleep), memory for emotionally toned information (negative, positive) decreased. No such effect was observed for neutral information.

**Figure 7.**
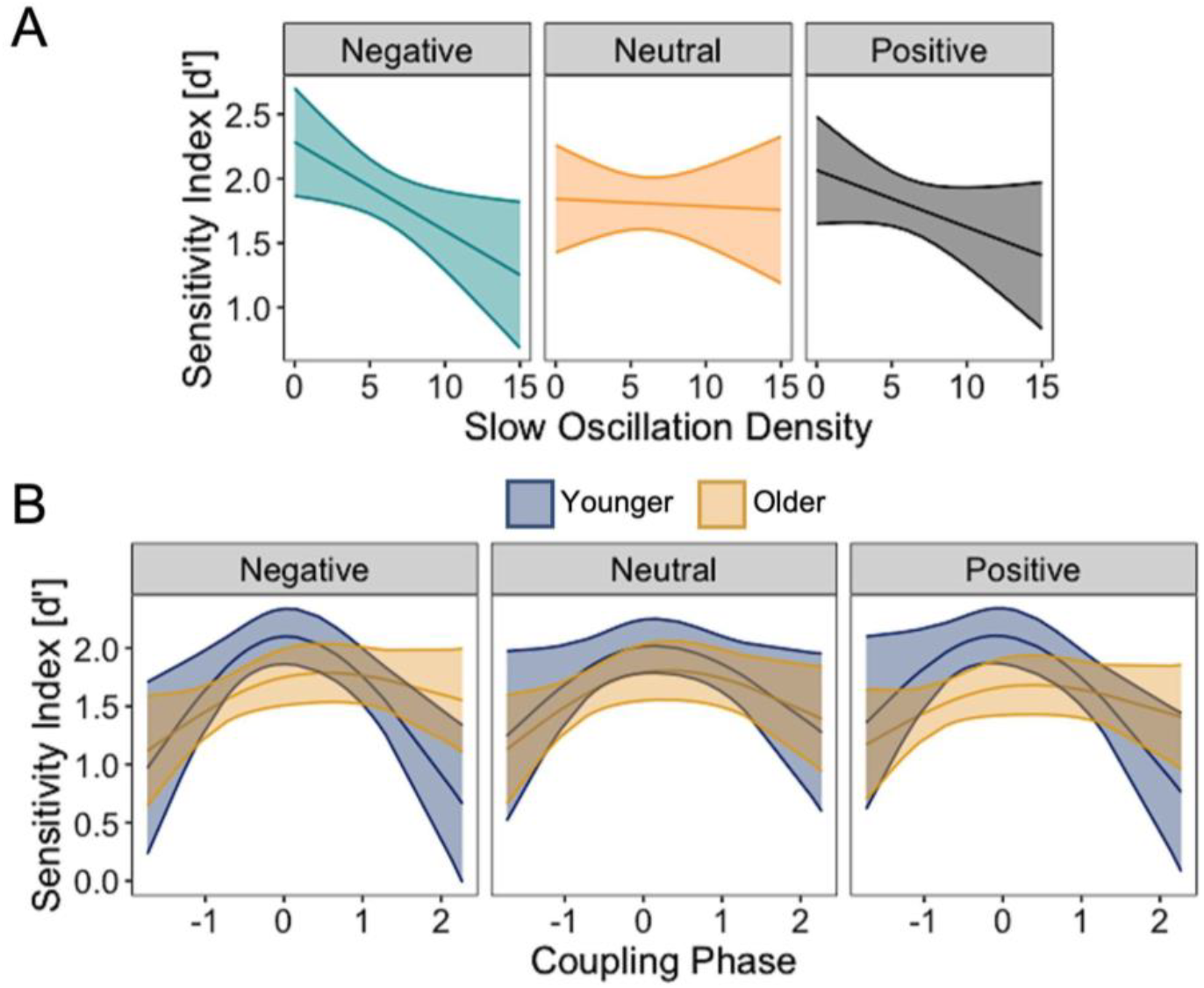
**(A)** Estimated marginal means for d’ scores at delayed testing by NREM SO density (x-axis; higher values denote greater SO events per 30s epoch during NREM sleep) and valence (negative = left, neutral = middle, positive = right). Shaded regions indicate the 83% CI. (B)

We also revealed a significant SO-Sigma Phase × Age × Valence interaction on recognition accuracy (χ2(4) = 18.17, *p* = .001; Figure 7B). Here, when sigma power was maximal closest to the peak of the SO (0 on the axis of Figure 7B), memory was strongest for all valences; however, this effect was largest for negative information and younger individuals, and weakest for neutral information and older individuals. This effect demonstrates that memory is strongest when sigma power peaked near the SO peak, with the greatest enhancement for negative information in younger individuals and the weakest for neutral information in older individuals.

## 5. Discussion

The current study examined how emotional memory consolidation varies across the lifespan as a function of interindividual differences in neural noise, quantified as the aperiodic slope, and how this relationship relates to sleep neurophysiology. Our results reveal significant differences in both emotional memory and aperiodic neural activity across age, highlighting complex interactions between neural processing, sleep, and memory consolidation. Specifically, we found that aperiodic activity modulates memory consolidation differentially across age, particularly for negative emotional content. These findings contribute to the growing body of literature suggesting that aperiodic neural activity, a measure of intrinsic neural “noise” (Cross et al., 2024; Voytek et al., 2015), can serve as a marker of neural efficiency, influencing emotional memory processes.

First, we revealed a nonlinear trajectory of memory performance across the lifespan, with performance peaking in young adulthood and declining thereafter. This is consistent with previous studies that describe an inverted u-shape in memory performance across the lifespan (e.g., Davis et al., 2003). Notably, we observed no main effect of emotional valence on recognition accuracy. This is in contrast to earlier research that has reported superior memory for negative stimuli (e.g., Kensinger & Schacter, 2006). Our results suggest that individual differences in brain activity, such as those captured by aperiodic neural activity, may be a stronger predictor of memory performance than emotional content alone. This finding supports the idea that memory is influenced not only by the emotional valence of the stimuli (Alger et al., 2018; Kensinger, 2009) but also by the brain’s intrinsic neural dynamics. We also found that aperiodic slopes followed a u-shaped trajectory across age, with steep slopes observed in childhood and older adulthood, and flatter slopes in middle age. These results align with previous studies suggesting that neural efficiency, as reflected by the aperiodic slope, varies across the lifespan (Merkin et al., 2023), and recent intracranial EEG work reporting nonlinear aperiodic dynamics across age (Cross et al., 2024). In younger adults, steeper aperiodic slopes are associated with more efficient brain processing (Voytek et al., 2015), potentially facilitating the consolidation of emotional memories. The flattening of slopes across age may reflect less efficient neural processing (Bornkessel-Schlesewsky et al., 2022; Tran et al., 2020), which could impair the brain’s ability to consolidate emotionally significant information. This suggests that age-related differences in aperiodic activity, which are indicative of neural efficiency, might contribute to differences in emotional memory consolidation across age.

Regarding the modulation of memory by sleep and wake states, we found that sleep enhances memory for negative stimuli, consistent with prior studies that highlight the beneficial role of sleep in emotional memory consolidation (Alger et al., 2019; Payne et al., 2008). More importantly, our results showed that the sleep-related enhancement of memory was influenced by interindividual differences in aperiodic slopes. Participants with steeper slopes exhibited better memory for negative stimuli following sleep, while those with flatter slopes showed a decline in memory performance with age. These findings extend previous work suggesting that individual differences in aperiodic activity may modulate sleep-dependent memory processes (Helfrich et al., 2021). Our results imply that individuals with more efficient neural processing (steeper slopes) may benefit more from sleep-based memory consolidation mechanisms, resulting in better retention of emotional memories.

We also observed that NREM SOs were associated with poorer memory for emotional information but not neutral content. This negative relationship suggests that excessive SO activity during sleep may hinder emotional memory consolidation. While these findings contrast with studies that have found a positive relationship between SOs and memory consolidation, our results point to the possibility that excessive SO activity may disrupt emotional memory consolidation by limiting the delicate hierarchical synchronization of sleep oscillations, impairing the transition between network-level and local processing during sleep important for memory processing (Staresina et al., 2016).

Additionally, we found that SO-sigma phase coupling predicted memory performance, with the strongest memory enhancement observed when sigma power peaked near the SO. This coupling was most beneficial for negative stimuli in younger individuals, with a decline in this effect in older adults. These findings are consistent with research showing that efficient SO-sigma coupling is crucial for effective memory consolidation (Mölle et al., 2011). Neocortically originating SOs provide a temporal framework for the coordination of neural processes important for the consolidation of newly acquired information. The up-state of SOs enhances cortical excitability, aligning spindles within the sigma frequency range with hippocampal sharp-wave ripples and thereby facilitating hippocampal-cortical communication, and promoting the redistribution and stabilization of memory traces (Helfrich et al., 2018; Staresina, 2024; Weiner et al., 2024). The age-related attenuation of this effect suggests that changes in sleep microstructure and the neural mechanisms underlying oscillatory coupling may contribute to the decline in emotional memory consolidation observed in older adults (Helfrich et al., 2018).

Our results suggest that differences in both aperiodic activity and NREM oscillations across age contribute to variations in emotional memory consolidation. These findings support theoretical models that propose a dual-process mechanism where aperiodic activity facilitates the switching between network-level and local processing during sleep, while NREM oscillations, such as SOs and sigma activity, support hippocampal-neocortical interactions crucial for memory consolidation (Helfrich et al., 2021). Our findings extend these models by demonstrating how age-related differences in both components can disrupt the balance between these processes, impairing emotional memory consolidation, and particularly in older adults.

While our study provides valuable insights, there are several limitations to consider. The uneven age distribution in our sample may limit the generalizability of the findings, particularly for younger and older participants. Future studies should aim for a more even distribution of participants across age and consider longitudinal analysis of aperiodic activity to better capture the dynamics of aperiodic activity and memory over time. Furthermore, individual differences in emotional reactivity and regulation were not assessed, and these factors may moderate the relationship between memory consolidation and neural activity. Including measures of emotional regulation or personality traits could provide further insight into how emotional processing influences memory consolidation. Finally, daytime sleep paradigms provide a convenient control for time-of-day and circadian effects (e.g. Alger et al., 2018; Bennion et al., 2016; Cellini et al., 2016; Gerstner & Yin, 2010; Nishida et al., 2009; Payne et al., 2015). However, with reductions in the proportion of REM and consolidation period length compared to typical overnight sleep paradigms, future studies should consider whether similar results are found following overnight sleep. Specifically, some evidence exists that memory superiority for emotional items is more apparent after longer periods containing a full night of sleep (Lipinska et al., 2019; Pierce & Kensinger, 2011; Wagner et al., 2006).

Taken together, the current study highlights the complex role of aperiodic neural activity and NREM oscillations in emotional memory consolidation across age. The differences we observed in both neural activity and memory performance suggest that intrinsic “neural noise” may play a crucial role in modulating sleep-dependent memory processes. Understanding these interactions may provide novel insights into the neurophysiological mechanisms underpinning emotional memory and offer potential avenues for interventions aimed at mitigating cognitive decline in aging populations.

